# VGLL3 Links Pericyte Hypercontractility to Perivascular Fibrosis of the Cerebral Microcirculation, a Novel Vasculopathy Leading to Distinct Long-Term Cerebral Autoregulation Dysfunction After Subarachnoid Hemorrhage

**DOI:** 10.64898/2026.08.25.747162

**Authors:** Yongjin Zhang, Yuncong Li, Chong Li, Hanfu Yu, Hongji Deng, Jianyang Yu, Haoming Xia, Chen Yu, Yang Zhang, Zhiling Luo, Yuehong Dong, Xiangbin Pan, Fei Wang

## Abstract

**BACKGROUND:** Cerebral ischemia following subarachnoid hemorrhage (SAH) has traditionally been considered transient because functional alterations of the cerebral microcirculation are thought to be self-limiting. However, we identified a previously unrecognized vasculopathy, perivascular fibrosis of the cerebral microcirculation (PFCM), characterized by excessive type I collagen deposition after SAH. This study investigated the mechanisms underlying PFCM and its subsequent effects on cerebral hemodynamics.

**METHODS:** *In vivo* SAH was modeled in mice by autologous blood injection, whereas oxygenated hemoglobin (OxyHb) exposure was used to mimic SAH *in vitro*. Pericyte-deficient mice (*Pdgfrβ^+/-^*) and pericyte-specific vestigial-like family member 3 (VGLL3) conditional knockout mice (*Vgll3^ΔPC^*) were generated. Pericyte contractility was measured by nanoindentation and traction force microscopy. Molecular mechanisms were examined using Western blotting, immunofluorescence, CUT&Tag, RNA-seq, transmission electron microscopy, and molecular docking. PFCM, impaired dilation of the cerebral microcirculation, and cerebral autoregulation were assessed by two-photon imaging, transcranial Doppler with continuous blood pressure monitoring, super-resolution ultrasound imaging, and photoacoustic imaging.

**RESULTS:** After SAH, mice developed long-term cerebral autoregulation dysfunction marked by impaired dilation of the cerebral microcirculation, with the abnormality being most evident within the relatively lower blood pressure range. The marked reduction in PFCM in *Pdgfrβ^+/-^* mice indicated that pericytes were the principal cellular contributors. Mechanistically, OxyHb-induced cytoskeletal remodeling *in vitro* increased pericyte contractility and promoted nuclear translocation of SAH-upregulated VGLL3. This was followed by increased genomic occupancy, *Col1a1* transcriptional activation, and type I collagen deposition. Pericyte-specific VGLL3 knockout abolished PFCM and, consequently, significantly alleviated long-term cerebral autoregulation dysfunction.

**CONCLUSIONS:** Our findings identify PFCM mediated by pericytic VGLL3 as a novel vasculopathy leading to long-term cerebral autoregulation dysfunction after SAH.

**Clinical Perspective:** *What Is New?:*

- PFCM is a previously unrecognized vasculopathy that is mediated by pericytic VGLL3 after SAH in mice.
- PFCM is associated with persistent cerebral autoregulation dysfunction and impaired dilation of the cerebral microcirculation, challenging the traditional concept that cerebral ischemia after SAH is only transient and self-limiting.
- Pericyte-specific VGLL3 knockout abolished PFCM and, consequently, significantly attenuated long-term cerebral autoregulation dysfunction after SAH.

*What Are the Clinical Implications?:*

- Suppressing pericyte hypercontractility reduced early cerebral ischemia and was accompanied by less PFCM and reduced persistent cerebral autoregulation dysfunction after SAH.
- Pericytic VGLL3 lies within the hypercontractility-PFCM pathway and may therefore represent a specific therapeutic target for hemodynamic impairment after SAH.
- The findings suggest that cerebral ischemia after SAH may be aggravated by hypovolemia or hypotension. This observation supports careful cardiovascular management and further evaluation of integrated heart-brain treatment strategies

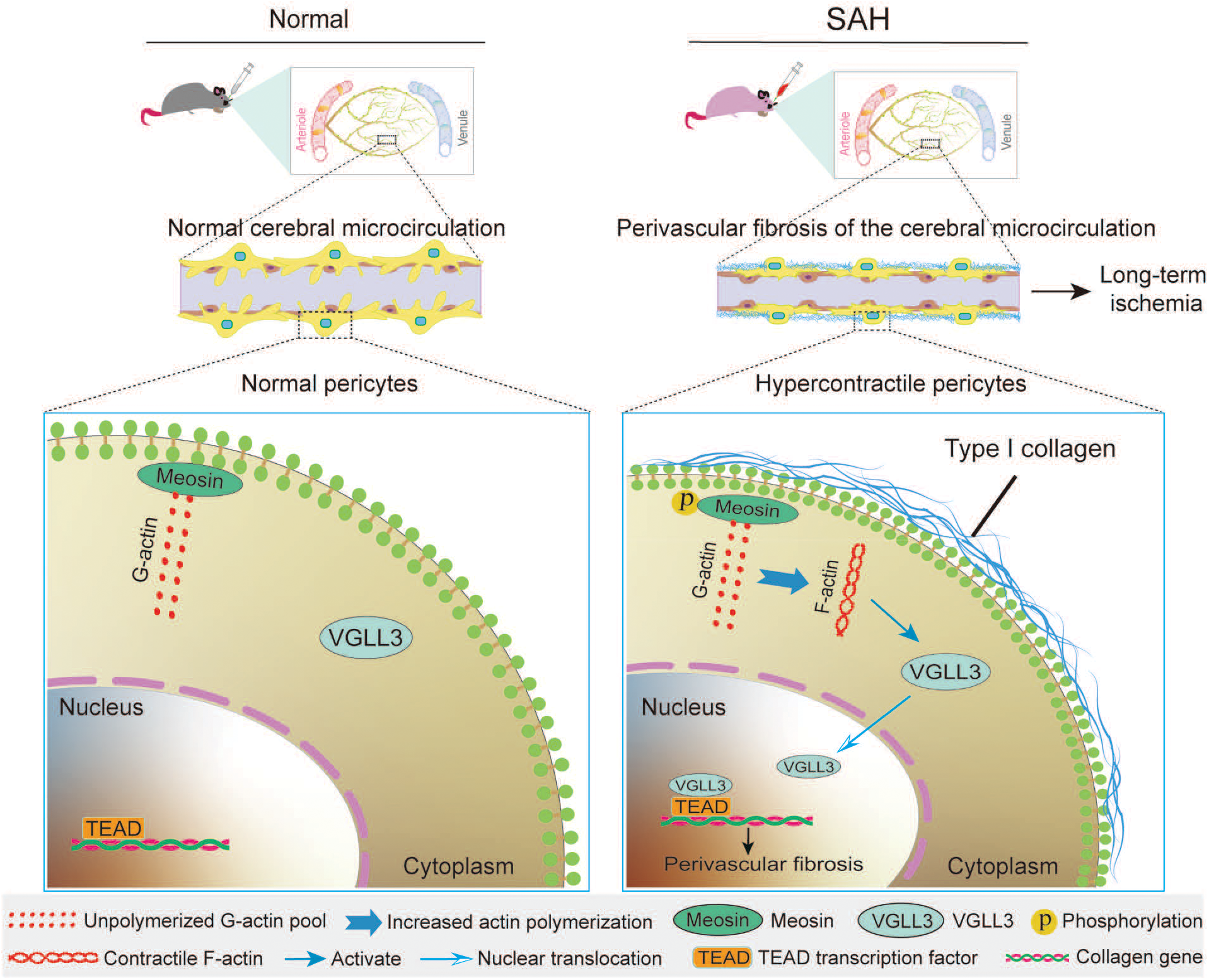

## INTRODUCTION

Subarachnoid hemorrhage (SAH) is a devastating cerebrovascular disorder primarily caused by ruptured intracranial aneurysms, associated with substantial long-term morbidity and mortality.^1,2^ Delayed cerebral ischemia (DCI) affects approximately 30% of SAH patients, typically occurring between 3 and 14 days after ictus.^3,4^ The pathophysiology of DCI has been extensively investigated, with impaired cerebral autoregulation recognized as a key contributing factor,^5,6^ and functional alterations of the cerebral microcirculation have been reported to contribute to transient ischemia after SAH.^7,8^

Accordingly, current clinical interventions largely focus on acute-phase vasospasm.^9^ This emphasis, however, does not fully account for evidence that ischemic consequences of SAH may persist. Neurological deficits have been documented in SAH survivors,^2,10^ and population-based cohort studies report an increased risk of recurrent ischemic events for up to 6 years after SAH.^11,12^ Sustained neurovascular uncoupling has also been observed 30 days after SAH in mice.^13^ These observations suggest that ischemic injury can extend beyond the acute DCI period. Because structural remodeling of the cerebral microcirculation contributes to chronic ischemia,^7,14^ it remains important to determine whether the brain undergoes perivascular fibrotic remodeling after SAH, a process well recognized in cardiac disease.^15^

To address this question, we first examined whether perivascular fibrosis occurs in the cerebral microcirculation after SAH. Pericytes are the primary contractile mural cells of the cerebral microcirculation^16^ and contract early after SAH.^17^ Accordingly, pericyte contractility orchestrates microcirculatory perfusion homeostasis under both physiological and pathological conditions.^18^ Excessive pericyte contraction generates pathological mechanical tension, which activates cytoskeleton-dependent mechanotransduction signals^19–21^ and downstream transcriptional reprogramming.^22,23^ However, the molecular mechanism linking pericyte contraction to mechanotransduction in SAH remains poorly understood. Notably, VGLL3, an evolutionarily conserved mechanosensitive cofactor, competes with YAP/TAZ for TEAD binding and translates extracellular mechanical cues into profibrotic transcriptional programs across diverse tissues.^24,25^ This makes VGLL3 a compelling candidate for mediating pericyte hypercontractility; however, its role in SAH remains unknown.

In this study, we investigated whether persistent perivascular fibrosis accompanies long-term cerebral autoregulation dysfunction in mice after SAH. Pericyte-deficient PDGFRβ heterozygous knockout mice were used to assess whether pericytes are the major cellular source driving PFCM. Because publicly available human SAH transcriptomic data are lacking, human brain single-cell transcriptomic datasets from ICH patients were used to prioritize pericyte-enriched genes responsive to hemorrhagic injury; VGLL3 emerged as a leading candidate for subsequent validation in the SAH setting. We then tested whether pericyte-specific *Vgll3* knockout could prevent PFCM and improve the accompanying autoregulatory dysfunction.

## METHODS

The corresponding author will provide the data, analytical methods, and study materials upon reasonable request.

### Animals

Approval for all animal procedures was obtained from the Animal Welfare and Ethics Committee of the First Affiliated Hospital of Kunming Medical University (Approval No. 2026DF041), and the procedures followed the NIH Guide for the Care and Use of Laboratory Animals. Animals were maintained at 23–26°C under a 12-h light/dark cycle with ad libitum access to food and water. Assignment to experimental groups was random, and each mouse was treated as an independent experimental unit.

Cas9-mediated microinjection into C57BL/6J zygotes was used to generate *Pdgfrβ^+/-^* mice (strain NM-KO-242464, Shanghai Model Organisms). F0 founders carrying confirmed mutations were backcrossed with C57BL/6J mice to obtain F1 heterozygotes. Because *Pdgfrβ^+/-^* homozygotes are embryonic lethal and exhibit severe cardiovascular, renal, and hematologic defects,^26^ only heterozygotes were studied; littermate *Pdgfrβ^+/+^* mice served as controls.

*Vgll3^ΔPC^* mice were produced by crossing *Vgll3^flox/flox^* (*Vgll3^fl/fl^*) (NM-CKO-241534) mice with *Pdgfrβ*-CreERT2 transgenic mice (NMX-TG-241616), both obtained from Shanghai Model Organisms. Cre-negative littermate *Vgll3^fl/fl^* mice served as controls. Genotyping primers are listed in the Major Resources Table. Cre recombination was induced in 10-week-old mice by tamoxifen administration (Sigma-Aldrich, 75 mg/kg/day, i.p.) for 5 consecutive days. Both *Vgll3^ΔPC^* mice and control littermates subsequently underwent SAH surgery, and brain tissues were harvested 7 days after the final tamoxifen injection.

### SAH animal model

SAH was induced as previously described.^27^ Briefly, mice were anesthetized with 2% isoflurane in 100% O₂, and anesthesia was maintained with 1% isoflurane. The mice were placed in a stereotaxic frame, and a midline skin incision was made over the skull. A burr hole was drilled 4.5 mm anterior to the bregma using a 0.9-mm drill until the dura was penetrated. A 27-gauge needle was advanced ventrally at a 40° angle through the burr hole to a depth of 4.5 mm below the dura. Then, 50 μL of autologous arterial blood was injected into the prechiasmatic cistern over 10 seconds. The needle was left in place for 3 minutes and then slowly removed to prevent backflow. The burr hole was immediately sealed with bone wax, and the incision was sutured. Mice were allowed to recover on a heating pad until fully awake before being returned to their cages. Sham-operated mice underwent the same procedures without blood injection. All surgical procedures and subsequent analyses were performed by investigators blinded to genotype and group allocation.

### Cerebral microcirculation, pericytes, and PFCM imaging

Two-photon microscopy was employed for real-time imaging of cerebral microcirculation and pericytes, with concurrent visualization of PFCM, as described previously.^28^ Acquired images were analyzed using ImageJ to quantify type I collagen deposition.

### Photoacoustic imaging

Cortical microvascular density and vessel tortuosity were assessed by photoacoustic microscopy using a GAni+Plus imaging system (Guangying Cell, China). Label-free signals were acquired at 532 nm and 1064 nm. Vascular density was quantified using ImageJ and expressed as the percentage of vascular area, and tortuosity as the ratio of actual vessel length to straight-line distance. Three to five randomly selected fields were analyzed per mouse.

### Transcranial Doppler with Continuous Blood Pressure Monitoring for Cerebral Autoregulation

Cerebral autoregulation was determined by continuous monitoring of cerebral blood flow velocity (CBFV) using a transcranial Doppler probe (20 MHz) fixed over the right parietal bone, with simultaneous invasive mean arterial pressure (MAP) recording via femoral artery catheterization. MAP was systematically manipulated across 40–190 mmHg by graded hemorrhage and phenylephrine infusion. At each pressure level, CBFV was recorded after a 3-minute equilibration period. For each mouse, CBFV was plotted against MAP, and a sigmoidal curve was fitted to determine the autoregulation plateau (CBFV within 20% of baseline) and the lower and upper limits of autoregulation. All analyses were performed blinded to genotype.

### Super-Resolution Ultrasound Imaging of Cerebral Microcirculation

Mice were anesthetized with 2% isoflurane and placed in a stereotactic frame. After shaving and aseptic preparation, a scalp incision was made and the periosteum retracted. A cranial window was made over the parietal cortex, and the headplate was secured. Ultrasound gel was instilled, and the probe (X10-23L) was positioned over the target region. Contrast agent was infused at 3.5 mL/h, and imaging was performed in contrast mode at 6,000–10,000 frames per session.

### Primary Pericyte Culture

Primary brain pericytes were isolated as previously described.^17^ Briefly, mouse brain tissue was digested enzymatically with papain and DNase I at 37°C for 60 minutes to generate pericyte-enriched cell suspensions. The suspensions were centrifuged at 3000×g for 5 minutes at 4°C, the supernatant was removed, and the cell pellet was resuspended in pericyte-specific medium. Cells were maintained at 37°C in a humidified 5% CO₂ atmosphere. Pericyte identity was verified by immunofluorescence (IF) staining for PDGFRβ and NG2 flow cytometry confirmed a purity of >85% before further culture and passage.

### VGLL3 Translocation Assay

For nuclear translocation analysis, cells on coverslips were pretreated with blebbistatin (10 μM, 30 min) followed by OxyHb (10 μM) for 12 h, or treated with OxyHb alone for 12, 24, or 72 h. After fixation and permeabilization, cells were immunostained for VGLL3 and counterstained with DAPI. Nuclear VGLL3 intensity per nuclear area was quantified using ImageJ.

### Type I collagen Deposition Assay

Primary brain pericytes were seeded onto Matrigel-coated coverslips and, after 12 hours, treated with OxyHb (10 μM) with or without inhibitors for 12, 24, or 72 hours. At each time point, cells were fixed and immunostained for type I collagen, with nuclei counterstained with DAPI. Images were captured by confocal microscopy, and the collagen-positive area was quantified using ImageJ and normalized to the number of nuclei (DAPI-positive cells).

### F/G-Actin Ratio Assay

The actin-to-G-actin ratio was determined using a commercial kit (Cytoskeleton, CSK-BK037). Cells were lysed in F-actin stabilizer buffer, and lysates were ultracentrifuged at 100,000×g for 1 hour at 4°C to separate F-actin (pellet) from G-actin (supernatant). Fractions were resolved by SDS-PAGE, band densities were quantified with ImageJ, and the F/G-actin ratio was calculated.

### Nanoindentation and traction force microscopy assessment of pericyte contractility

Pericytes exposed to OxyHb (10 μM) were evaluated by nanoindentation and traction force microscopy (TFM). Nanoindentation used a Piuma nanoindenter (Hertz Optics) equipped with a Berkovich tip (max load 98 nN; loading rate 10 nm/s). Elastic modulus and hardness were obtained from loading/unloading curves using the Oliver–Pharr method. Stress relaxation was determined from the hold segment at peak load, and the percentage of relaxation over time was calculated. For TFM, pericytes were grown on bead-embedded polyacrylamide gels in a XuanYuan TFM system, and traction stress was derived by Fourier transform traction cytometry. OriginPro (10.3) was used for data analysis. At least 30 cells per condition were analyzed.

### Cerebral Microvascular Dilation Assay

Cerebral microvessels in living brain slices were examined by infrared differential interference contrast (IR-DIC) two-photon microscopy with a 40× water-immersion objective. Lumen diameter was measured before and after bath exposure to Ca²⁺-free and 2 mM Ca²⁺-containing solutions. For each vessel segment, measurements from three sites were averaged to calculate the maximum dilation ratio: [(Dpassive − Dactive) / Dpassive] × 100%, where Dpassive and Dactive represent diameters in Ca²⁺-free and 2 mM Ca²⁺ solutions, respectively. The calculated ratio reflects microvascular wall compliance and maximal dilation capacity.^29^

### CUT&Tag and RNA-seq

For CUT&Tag, libraries were sequenced on NovaSeq 6000 (PE150). Reads were trimmed with Trimmomatic (v0.39), and peaks were called using MACS2 (v2.2.7.1). VGLL3 occupancy at promoter regions was analyzed, and GO/KEGG enrichment was performed with ClusterProfiler (v4.0).

For RNA-seq, total RNA was extracted and sequenced on a NovaSeq 6000 platform. The obtained reads were aligned to the mouse reference genome using HISAT2 (v2.1.0), and differentially expressed genes (DEGs) were defined as |log₂FC| > 1, and adjusted *p*-value (*p*adj) < 0.05. GO and KEGG enrichment analyses were performed using ClusterProfiler (v4.0).

### Statistical analysis

Data were analyzed and graphs were generated using Prism 8 (GraphPad Software). All data are presented as individual data points with mean ± standard error of the mean (SEM). For comparisons between two groups, the Student’s t-test (two-tailed) was used, with Welch’s correction applied when variances were unequal. For comparisons among three or more groups, one-way ANOVA was performed, followed by Tukey’s post hoc test for multiple comparisons. A two-tailed *p* < 0.05 was considered statistically significant. Significance levels are indicated as follows: \**p* < 0.05, \*\**p* < 0.01, \*\*\**p* < 0.001, and \*\*\*\**p* < 0.0001; n.s., not significant. The number of independent experiments (n) is specified in each figure legend.

## RESULTS

### SAH induces PFCM and persistent cerebral autoregulation dysfunction with impaired dilation of the cerebral microcirculation

Studies in cardiac disease have identified perivascular fibrosis of the microcirculation as a key pathological feature, characterized by excessive perivascular collagen deposition.^30^ To determine whether a similar perivascular remodeling occurs in the brain after SAH, we examined the cerebral microcirculation and observed a progressive widening of the perivascular space after SAH (Figure 1A and 1G). Concurrently, extensive type I collagen deposition was detected around the cerebral microcirculation in SAH mice at 1, 7, and 30 days after SAH, indicative of progressive PFCM (Figures 1B, 1H, and S1A, S1B). Photoacoustic imaging further revealed increased cerebrovascular tortuosity and reduced vascular branches at 7 and 30 days after SAH, along with a moderate decrease in cerebral microvascular density (Figures 1C, 1I, 1J, and S1C, S1D). Collectively, these findings demonstrate that SAH induces progressive PFCM and microvascular rarefaction, prompted by structural alterations. Next, we investigated whether PFCM translates into functional impairment, to examine whether these structural alterations translate into functional impairment of the cerebral microcirculation.

**Figure 1.**
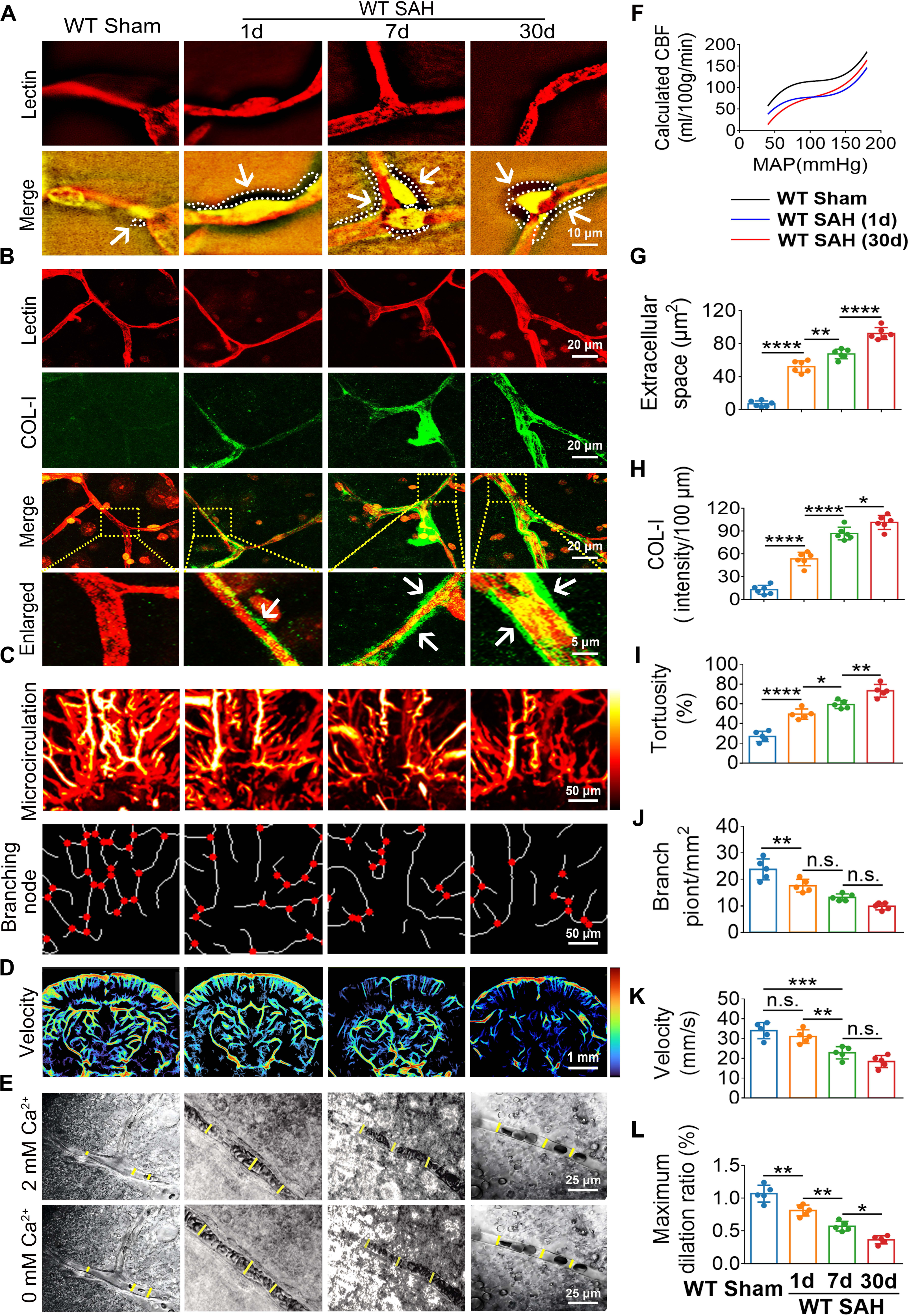
PFCM and persistent cerebral autoregulatory dysfunction after SAH. **A, G,** N-SIM super-resolution images and measurements of the perivascular space surrounding cerebral pericytes in C57BL/6J mice at the indicated time points after SAH. Scale bar, 10 μm. **B, H,** Ex vivo brain-slice images showing perivascular type I collagen (COL-I, green) around the cerebral microcirculation (Lectin, red), with pericytes labeled by TO-PRO-3 (orange). Scale bar, 20 μm. **C, I, and J,** Photoacoustic measurements of cerebrovascular tortuosity and vascular branch nodes. After SAH, tortuosity increased progressively while the number of branch nodes declined, and both changes were greatest at 30 days after SAH. Scale bar, 50 μm. **D, K,** Mean cerebral blood flow velocity assessed by super-resolution ultrasound imaging at 1, 7, and 30 days after SAH. A progressive reduction was detected relative to sham controls (MAP ≈ 60 mmHg). **E, L,** Vasodilatory capacity of the cerebral microcirculation evaluated after application of Ca²⁺-free artificial cerebrospinal fluid (ACSF). Dilation was impaired after SAH, with the largest deficit at 30 days after SAH. **F,** Cerebral autoregulation curves derived from cerebral blood flow velocity and mean arterial pressure (MAP). At 1 day after SAH, the curve showed a global downward shift (blue curve); at 30 days after SAH, the downward shift was concentrated mainly in the lower blood pressure range (MAP < 80 mmHg, red curve). Statistical analysis: Data are presented as mean ± SEM (n = 6 mice per group). Two-group comparisons were performed using unpaired, two-tailed Student’s t-test. Comparisons among three or more groups were performed by one-way ANOVA followed by Tukey’s post hoc test. \**p* < 0.05, \*\**p* < 0.01, \*\*\**p* < 0.001, \*\*\*\**p* < 0.0001. Abbreviations: PFCM, perivascular fibrosis of the cerebral microcirculation; SAH, subarachnoid hemorrhage; COL-I, type I collagen; ACSF, artificial cerebrospinal fluid; MAP, mean arterial pressure; N-SIM, non-linear structured illumination microscopy; SEM, standard error of the mean; n.s., not significant.

DCI is commonly associated with impaired cerebral autoregulation and reduced vasodilatory capacity of the cerebral microcirculation.^6,31^ Cerebral autoregulation curves were therefore generated by measuring cerebral blood flow velocity over a range of blood pressures (Figure 1F). At 1 day after SAH, the curve showed a global downward shift (Figure 1F, blue curve), consistent with an early contribution from cerebral microcirculatory vasospasm. By 30 days after SAH, the downward shift was largely restricted to the lower blood pressure range (Figure 1F, red curve). Super-resolution ultrasound imaging supported this pattern by showing a progressive reduction in cerebral blood flow velocity at 1, 7, and 30 days after SAH (Figures 1D and 1K). Cerebral microvessel diameter was then measured to examine vasodilatory capacity (Figure 1E). Perfusion with Ca²⁺-free artificial cerebrospinal fluid (ACSF) revealed significantly impaired microvascular dilation after SAH (Figures 1E and 1L), suggesting that impaired dilation may contribute substantially to long-term (30-day) cerebral ischemia after SAH.

Taken together, our data suggest that SAH induces PFCM and long-term cerebral autoregulation dysfunction, characterized by impaired dilation of the cerebral microcirculation, particularly at lower blood pressures.

### Pericytes are principally responsible for PFCM after SAH in mice

We first confirmed pericyte distribution along the cerebral microcirculation, with TO-PRO-3 pericytes closely apposed to the Lectin endothelial tube (Figure 2B). To determine whether pericytes mediate PFCM, we evaluated collagen deposition using both *in vitro* and *in vivo* SAH models. *In vitro*, primary pericytes were treated with OxyHb to mimic SAH (Figure 2A), which induced time-dependent type I collagen deposition (Figures 2C, 2D, and 2F).

**Figure 2.**
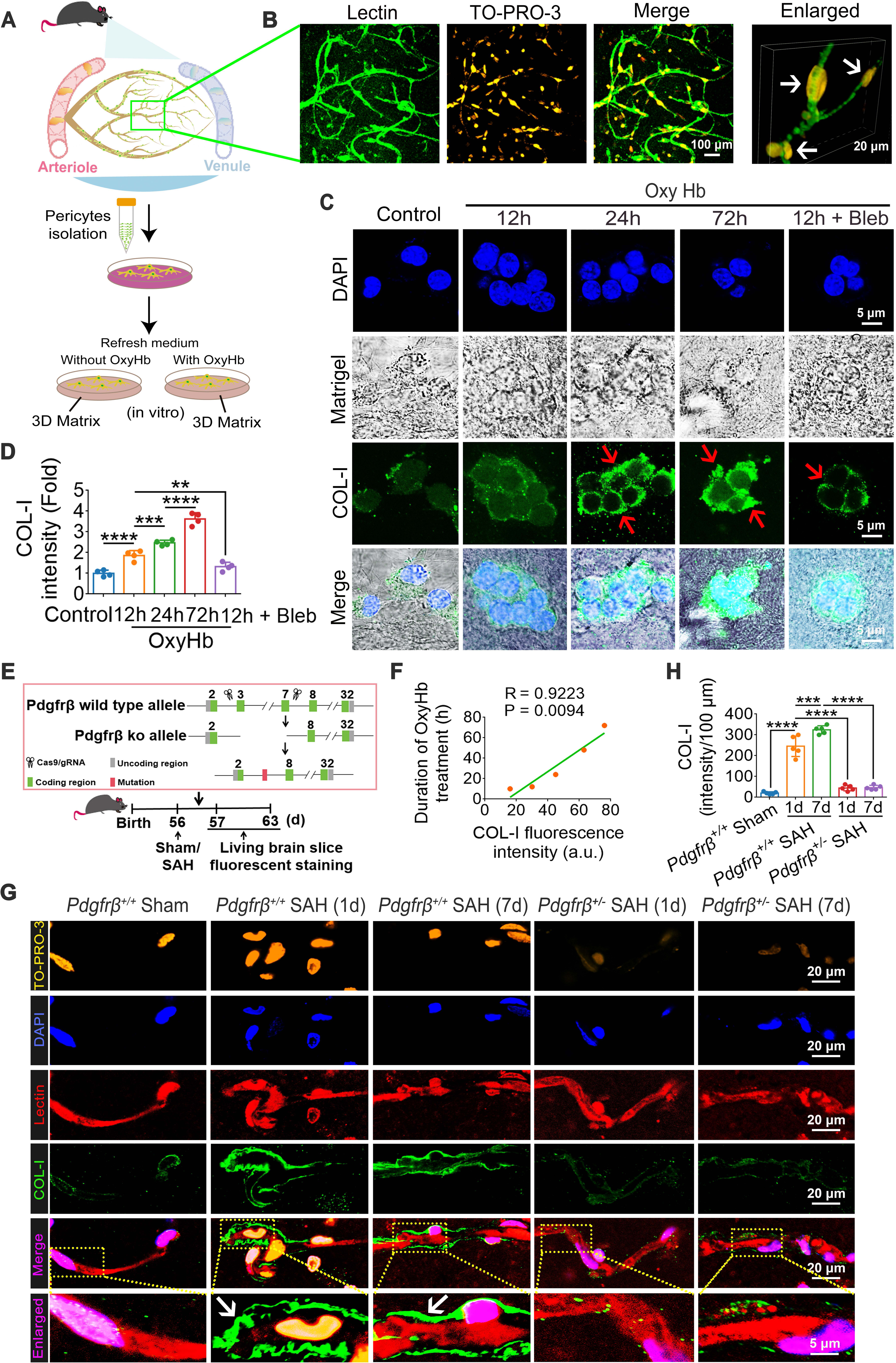
Pericytes are the principal contributors to PFCM after SAH in mice. **A,** Schematic of primary mouse brain pericyte isolation followed by OxyHb exposure to reproduce the SAH microenvironment *in vitro*. **B,** Representative IF images of pericytes (TO-PRO-3, orange) and the cerebral microcirculation (Lectin, green) in *ex vivo* brain slices from *Pdgfrβ*⁺/⁺ littermate mice. Pericytes lie along the microvascular wall, with elongated cell bodies and branched processes surrounding the underlying endothelium. Scale bars: 100 μm (overview) and 20 μm (enlarged view). **C, D,** Representative IF images and corresponding quantification of type I collagen (COL-I, green) in primary pericytes exposed to OxyHb with or without Bleb, a non-muscle myosin II inhibitor of pericyte contraction. OxyHb produced a time-dependent increase in collagen deposition, whereas Bleb co-treatment significantly reduced this response. n = 4 independent experiments. Scale bar, 5 μm. **E,** Experimental scheme for *in vivo* validation in *Pdgfrβ*⁺/⁻ mice. **F,** Quantification of type I collagen deposition after 6, 12, and 24 hours of OxyHb exposure, demonstrating progressive collagen accumulation (n = 4 independent experiments). Data are presented as mean ± SEM. \**p* < 0.05, \*\**p* < 0.01 vs. control; one-way ANOVA followed by Tukey’s post hoc test. **G, H,** Representative IF images and quantification of perivascular type I collagen (COL-I, green) surrounding the cerebral microcirculation (Lectin, red) in *Pdgfrβ^+/+^* sham, SAH-*Pdgfrβ^+/+^*, and SAH-*Pdgfrβ^+/-^* mice at 1 and 7 days after SAH. Perivascular collagen deposition after SAH was significantly lower in pericyte-deficient mice. Scale bar, 20 μm. n = 6 mice per group. Data are presented as mean ± SEM. \*\*\**p* < 0.001, \*\*\*\**p* < 0.0001; n.s., not significant; one-way ANOVA followed by Tukey’s post hoc test Abbreviations: PFCM, perivascular fibrosis of the cerebral microcirculation; SAH, subarachnoid hemorrhage; OxyHb, oxygenated hemoglobin; COL-I, type I collagen; Bleb, blebbistatin; SEM, standard error of the mean; n.s., not significant.

To validate these findings *in vivo*, *Pdgfrβ^+/-^* and *Pdgfrβ^+/+^* mice were subjected to SAH (Figure 2E). Genotyping confirmed the expected *Pdgfrβ* alleles (Figure S2A). Live brain slice staining with Lectin (cerebral microcirculation) and TO-PRO-3 (pericytes) revealed a significant reduction in pericyte coverage in *Pdgfrβ^+/-^* mice compared with *Pdgfrβ^+/+^* littermates (Figures S2B and S2C), confirming the successful reduction of pericyte abundance in the heterozygous mice. The progressive perivascular deposition of type I was significantly reduced in *Pdgfrβ^+/-^* mice compared with *Pdgfrβ^+/+^* littermates (Figures 2G and 2H). Collectively, these findings demonstrate that pericytes are the principal cells driving PFCM after SAH in mice.

Notably, we found that the OxyHb-induced type I collagen deposition *in vitro* could be significantly attenuated by Bleb, an inhibitor of cell contraction (Figure 2C). This result suggests that pericyte hypercontraction and mechanical tension may serve as upstream biomechanical triggers for their profibrotic phenotypic transition.

### OxyHb-induced cytoskeletal remodeling promotes pericyte hypercontractility, VGLL3 nuclear translocation, and subsequent type I collagen deposition *in vitro*

After determining pericytes as major contributors to PFCM, the molecular events underlying this response were examined *in vitro*. OxyHb was firstly tested for its impact on moesin phosphorylation, which regulates cytoskeletal dynamics and has been associated with fibrogenic gene expression.^32^ IF staining exhibited progressive increases in p-moesin and total moesin at 12, 24, and 72 hours after OxyHb stimulation (Figures 3A–3D), while WB independently verified time-dependent upregulation of p-moesin (Figures S3A and S3B). In addition, OxyHb also shifted actin from the G-actin pool toward F-actin (Figures 3E and 3F), confirming to enhanced polymerization. TFM measurements exhibited higher pericyte contractility after OxyHb treatment (Figure 3G). Then, nanoindentation was employed to characterize the biomechanical phenotype (Figure 3H, schematic adapted from Optics11; representative load-indentation curves presented in Figure S4A). Elastic modulus and hardness increased (Figure 3I), while stress relaxation was impaired (Figure 3J), suggesting greater stiffness and a hypercontractile state. Additionally, cell area also decreased progressively after OxyHb treatment (Figure 3O), offering an additional measure of the contractile phenotype.

**Figure 3.**
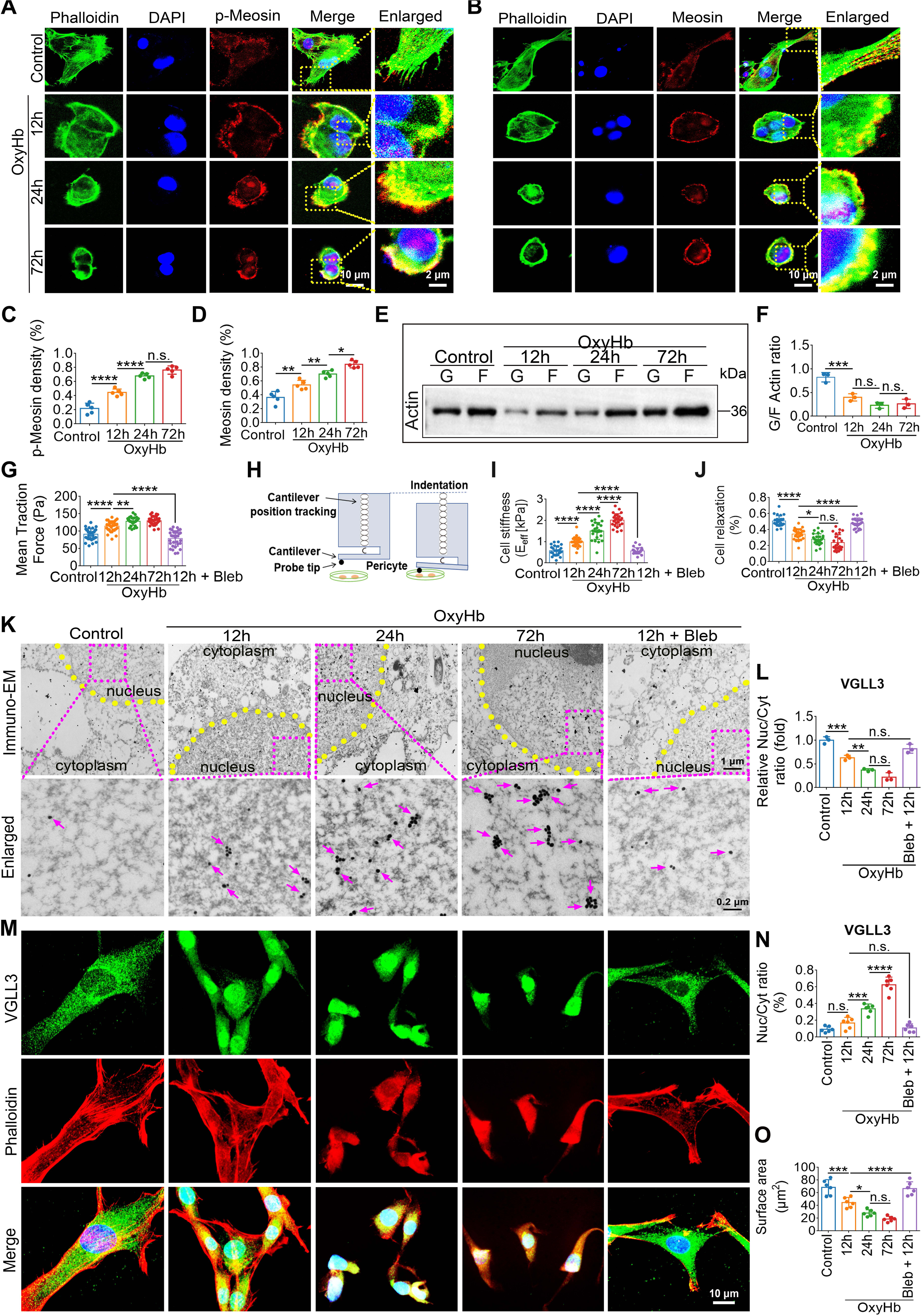
OxyHb-induced cytoskeletal remodeling promotes pericyte hypercontractility and nuclear translocation of SAH-upregulated VGLL3, followed by type I collagen deposition in vitro. **A, B, C, and D**, Representative IF images and measurements of p-moesin and total moesin in primary pericytes at 12, 24, and 72 hours after stimulation with OxyHb (10 μM). Both p-moesin and total moesin increased progressively over time after OxyHb exposure. Scale bars: 10 μm (overview) and 2 μm (enlarged view). n = 4 independent experiments. Data are indicated as mean ± SEM. \**p* < 0.05, \*\**p* < 0.01, \*\*\*\**p* < 0.0001 vs. control; one-way ANOVA followed by Tukey’s post hoc test. **E and F,** Representative WB images and quantification of the F-/G-actin ratio after OxyHb (10 μM) stimulation. A pronounced shift from G-actin toward F-actin was observed, consistent with enhanced actin polymerization. n = 4 independent experiments. Data are presented as mean ± SEM. \*\*\**p* < 0.001 vs. control; unpaired, two-tailed Student’s t-test. **G,** Traction force microscopy (TFM) measurement of pericyte contractility after OxyHb (10 μM) stimulation. Pericyte traction stress increased significantly after treatment. n = 30 cells per condition. Data are denoted as mean ± SEM. \*\*\**p* < 0.001 vs. control; unpaired, two-tailed Student’s t-test. **H,** Schematic of the nanoindenter probe before (left) and during (right) cell indentation, adapted from Optics11. **I, J,** Nanoindentation measurements in primary pericytes after OxyHb (10 μM) stimulation. Elastic modulus and hardness were obviously increased (I), while stress relaxation was impaired (J), indicating greater cellular stiffness and hypercontractility. n = 30 cells per condition. Data are presented as mean ± SEM. \**P* < 0.05, \*\*\*\**p* < 0.0001 vs. control; unpaired, two-tailed Student’s t-test. **K, L, M, and N,** Immuno-EM and IF assessment of VGLL3 nuclear translocation in primary pericytes after OxyHb (10 μM) stimulation. Nuclear VGLL3 signal increased progressively with time (K, M, L, N), and Immuno-EM verified localization of VGLL3 within the nucleus (K). Scale bars: 10 μm (overview) for IF; Scale bars: 1 μm (overview) and 200 nm (Enlarged) for Immuno-EM. n = 4 independent experiments. Data are presented as mean ± SEM. **P < 0.01, \*\*\**p* < 0.001 vs. control; one-way ANOVA followed by Tukey’s post hoc test. **O,** Cell-area quantification in primary pericytes after OxyHb (10 μM) stimulation. Cell area declined progressively over time, further supporting the contractile phenotype. n = 30 cells per condition. Data are presented as mean ± SEM. \**p* < 0.05, \*\**p* < 0.01 vs. control; one-way ANOVA followed by Tukey’s post hoc test. Abbreviations: OxyHb, oxygenated hemoglobin; p-moesin, phosphorylated moesin; TFM, traction force microscopy; Immuno-EM, immunoelectron microscopy; SEM, standard error of the mean; n.s., not significant.

Subsequently, we searched for transcriptional regulators that could relate OxyHb-evoked mechanical changes to profibrotic gene expression. Since no public human SAH transcriptomic dataset was available, the human ICH dataset GSE24265 was utilized as a hemorrhagic-injury surrogate. Differentially expressed genes were determined through |log₂FC| > 1 and adjusted *p*-value (*p*adj) < 0.05, and VGLL3 was strongly upregulated in ICH tissue (Figures S5A and S5B). Then, the human single-cell RNA-seq dataset GSE266873, comprising ICH samples from multiple time points and 10 resolved cell clusters (Figure S5C), was examined to determine the cellular distribution of VGLL3. VGLL3 expression was concentrated in the pericyte cluster (Figures S5D and S5E). On this basis, its response to OxyHb-induced hypercontractility was tested in primary pericytes. Immuno-EM and IF both revealed nuclear redistribution of VGLL3 after OxyHb stimulation (Figures 3K–3N), and WB revealed a progressive increase in VGLL3 protein expression (Figures S3C and S3D).

As shown in Figures 2C, 2D, and 2F, OxyHb stimulation led to a time-dependent increase in type I collagen deposition around pericytes. We next examined the functional relevance of VGLL3 in OxyHb-induced collagen deposition. Masson staining further confirmed progressive collagen accumulation up to 72 hours after OxyHb stimulation. Notably, VGLL3 knockdown markedly reduced collagen deposition at both 12 and 72 hours, whereas VGLL3 overexpression enhanced it, particularly at 72 hours (Figures S6A and S6B). These findings establish VGLL3 as a critical downstream effector of OxyHb-induced pericyte hypercontractility, driving profibrotic collagen deposition.

Having identified VGLL3 as a key regulator of collagen deposition in pericytes, we next investigated whether VGLL3 directly promotes collagen gene transcription. To this end, we performed CUT&Tag to map VGLL3 genomic occupancy.

### Pericyte-specific VGLL3 knockout attenuates PFCM after SAH in mice

To test whether VGLL3 is required for PFCM *in vivo*, we generated *Vgll3^ΔPC^* mice by crossing *Vgll3^fl/fl^* mice with *Pdgfrβ*-CreERT2 transgenic mice^33^ for tamoxifen-inducible pericyte-specific knockout (Figure 4A). At 10 weeks of age, mice were administered tamoxifen, then subjected to SAH, and brain tissues were collected at 1 and 7 days after SAH. IF staining revealed markedly reduced perivascular collagen I deposition in *Vgll3^ΔPC^* mice compared with *Vgll3^fl/fl^* littermates at both time points after SAH (Figure 4B and 4D).

**Figure 4.**
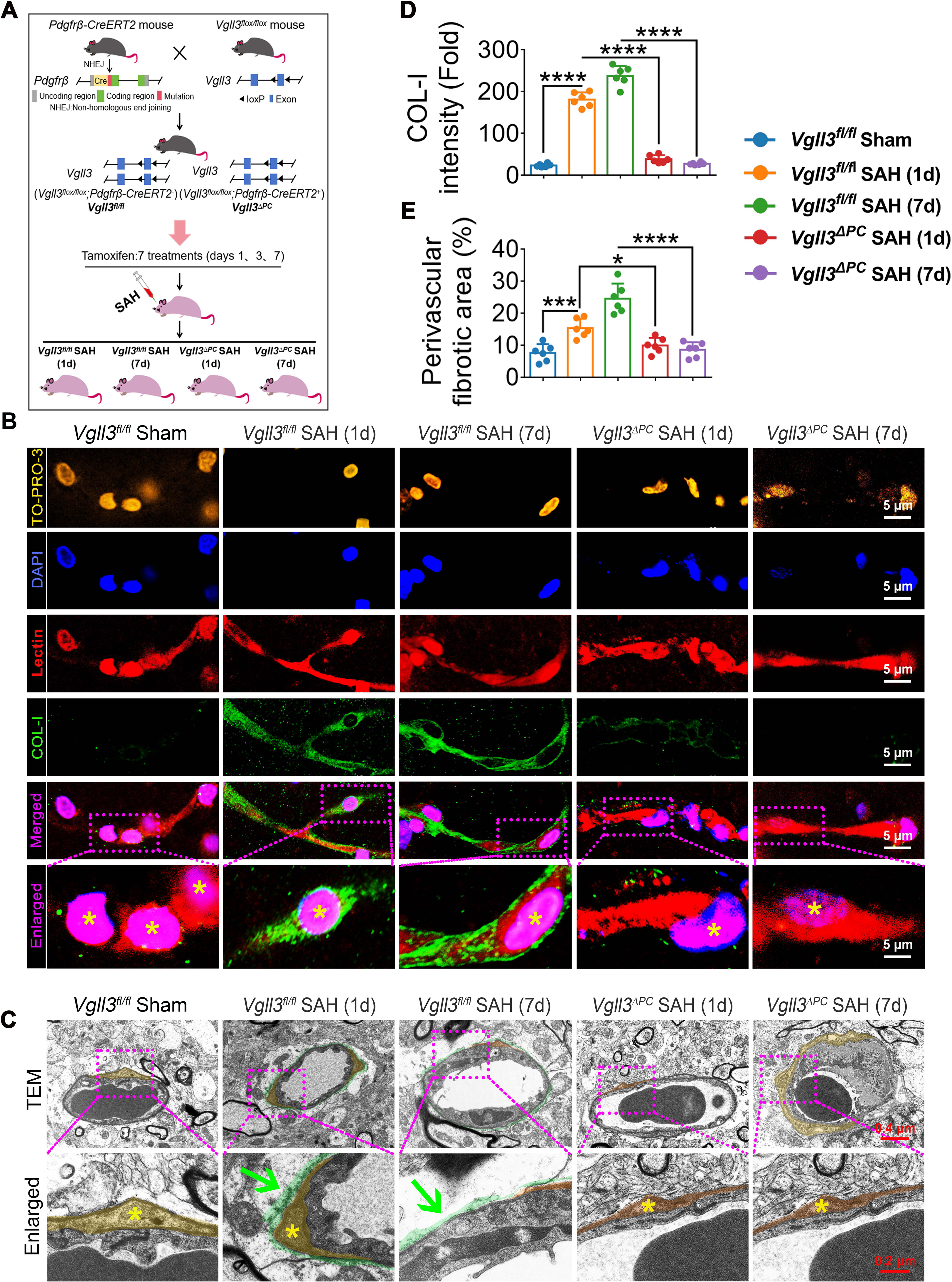
Pericyte-specific VGLL3 deletion reduces PFCM after SAH in mice. **A,** Schematic of the strategy used to generate pericyte-specific VGLL3 conditional knockout mice. *Vgll3^fl/fl^* mice were crossed with *Pdgfrβ*-CreERT2 transgenic mice; at 10 weeks of age, tamoxifen was administered to induce Cre-mediated VGLL3 deletion in pericytes before SAH induction. **B and D,** Representative IF images and quantification of perivascular type I collagen (COL-I, green) around the cerebral microcirculation (Lectin, red) in *Vgll3^fl/fl^* and *Vgll3^ΔPC^* littermate mice at 1 and 7 days after SAH. At both time points, perivascular collagen deposition was significantly lower after pericyte-specific VGLL3 knockout than in *Vgll3^fl/fl^* littermates. Scale bar, 5 μm. n = 6 mice per group. Data are presented as mean ± SEM. \*\*\**p* < 0.001, ****P < 0.0001; n.s., not significant; two-way ANOVA followed by Tukey’s post hoc test. **C,** Representative TEM images of perivascular collagen fibrils surrounding the cerebral microcirculation in *Vgll3^fl/fl^* and *Vgll3^ΔPC^* mice at 7 days after SAH. Ultrastructural collagen fibril accumulation was significantly lower in *Vgll3^ΔPC^* mice. Scale bar, 500 nm. n = 6 mice per group. **E,** Measurement of the perivascular fibrotic area in *Vgll3^fl/fl^* and *Vgll3^ΔPC^* mice at 1 and 7 days after SAH. *Vgll3^ΔPC^* mice showed a significant reduction in fibrotic area relative to *Vgll3^fl/fl^* littermates at both time points. n = 6 mice per group. Data are presented as mean ± SEM. \*\**p* < 0.01, ***P < 0.001; two-way ANOVA followed by Tukey’s post hoc test. All data are presented as mean ± SEM. \**p* < 0.05, \*\**p* < 0.01, \*\*\**p* < 0.001, \*\*\*\**p* < 0.0001. *Vgll3^fl/fl^* mice, *Vgll3flox/flox* mice; *Vgll3^ΔPC^* mice, pericyte-specific *Vgll3* conditional knockout mice. PFCM, perivascular fibrosis of cerebral microcirculation; SA**H,** subarachnoid hemorrhage; COL-**I,** type I collagen; TEM, transmission electron microscopy.

This was further supported by TEM analysis, which revealed a significant reduction in perivascular collagen fibril accumulation in *Vgll3^ΔPC^* mice relative to *Vgll3^fl/fl^* littermates at the ultrastructural level after SAH (Figure 4C). Quantitative analysis of the perivascular fibrotic area confirmed a significant decrease in the *Vgll3^ΔPC^* group (Figure 4E). Collectively, these findings demonstrate that pericytic VGLL3 is required for PFCM after SAH in mice.

### OxyHb promotes VGLL3 genomic occupancy and *Col1a1* transcriptional activation in pericytes *in vitro*

Given that pericytic VGLL3 is required for PFCM after SAH *in vivo* (Figure 4), we next investigated the molecular basis of VGLL3-mediated fibrogenic gene regulation. To this end, we performed CUT&Tag and RNA-seq on primary brain pericytes treated with OxyHb under *in vitro* conditions (Figure 5A).

**Figure 5.**
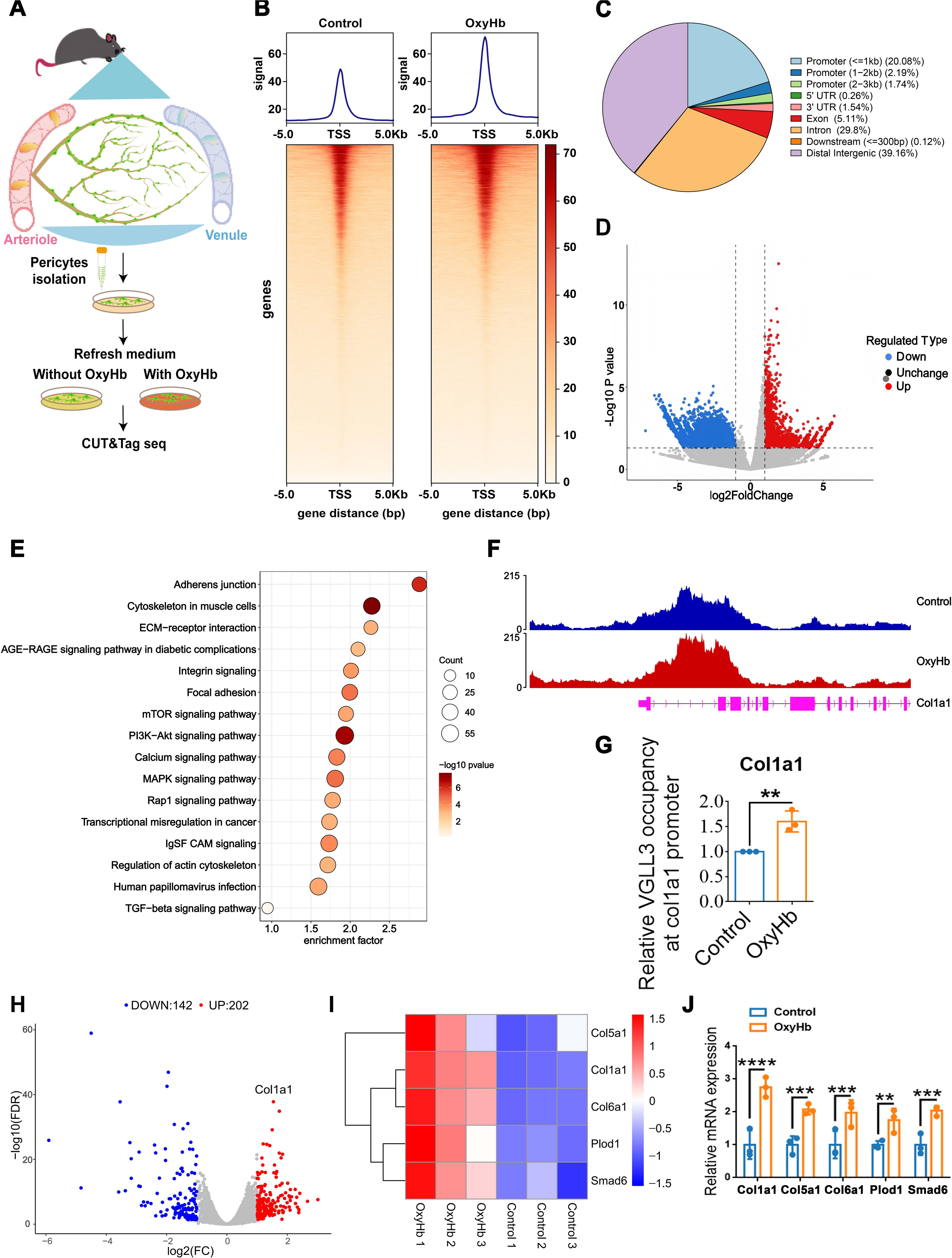
OxyHb increases VGLL3 genomic occupancy and activates *Col1a1* transcription in pericytes in vitro. **A,** Experimental scheme for CUT&Tag and RNA-seq in primary brain pericytes treated with OxyHb to model SAH *in vitro*. **B,** Metagene profile of VGLL3 CUT&Tag peaks relative to transcriptional start sites (TSS) in control and OxyHb-treated pericytes. OxyHb was associated with prominent VGLL3 enrichment within the 5-kb region flanking TSS. **C,** Distribution of VGLL3-occupied peaks in relation to TSS. Promoter-proximal regions accounted for 24.01% of all peaks (≤1 kb: 20.08%; 1–2 kb: 2.19%; 2–3 kb: 1.74%), indicating preferential occupancy near promoters. **D,** Differential VGLL3 CUT&Tag binding in control versus OxyHb-treated pericytes. OxyHb treatment was associated with 10,661 downregulated and 3,097 upregulated binding peaks, indicating broad redistribution of VGLL3 chromatin occupancy. **E,** KEGG enrichment analysis of genes adjacent to upregulated VGLL3 peaks. Matrix remodeling-related pathways were enriched, including PI3K-Akt signaling, ECM-receptor interaction, focal adhesion, and the TGF-β/Smad pathway. **F,** Genome browser tracks of VGLL3 CUT&Tag signal at the *Col1a1* locus in control and OxyHb-treated pericytes, demonstrating occupancy at the *Col1a1* promoter region. **G,** ChIP-qPCR assessment of VGLL3 occupancy at the *Col1a1* promoter in control and OxyHb-treated pericytes. OxyHb significantly increased VGLL3 enrichment at this promoter. Data are presented as mean ± SEM. \*\**p* < 0.01 vs. control; unpaired, two-tailed Student’s t-test. **H,** Volcano plot of RNA-seq data from OxyHb-treated pericytes and controls. In total, 202 genes were upregulated, and 142 genes were downregulated, with *Col1a1* showing the strongest induction. **I,** Heatmap of fibrosis-associated gene expression in control and OxyHb-treated pericytes. Expression of *Col1a1*, *Col5a1*, *Col6a1*, *Plod1*, and *Smad6* was significantly increased by OxyHb. **J,** RT-qPCR validation of fibrosis-associated gene expression in control and OxyHb-treated pericytes. Consistent with RNA-seq, RT-qPCR confirmed upregulation of *Col1a1*, *Col5a1*, *Col6a1*, *Plod1*, and *Smad6*. Data are presented as mean ± SEM. \**p* < 0.05, \*\**p* < 0.01, \*\*\**p* < 0.001, \*\*\*\**p* < 0.0001 vs. control; unpaired, two-tailed Student’s t-test. Abbreviations: OxyHb, oxygenated hemoglobin; TSS, transcriptional start site; KEGG, Kyoto Encyclopedia of Genes and Genomes; ECM, extracellular matrix; RT-qPCR, reverse transcription quantitative polymerase chain reaction; SEM, standard error of the mean; n.s., not significant.

We first mapped the genome-wide binding profiles of VGLL3 in pericytes exposed to SAH-mimicking conditions. Analysis of peak distribution relative to transcriptional start sites (TSS) revealed that OxyHb treatment induced prominent VGLL3 enrichment within the 5-kb flanking region of TSS (Figure 5B). Across all samples, 24.01% of VGLL3-occupied peaks were located in promoter-proximal regions, with the majority distributed within ≤1 kb upstream of TSS (Figure 5C). Differential binding analysis further identified 10,661 downregulated and 3,097 upregulated binding peaks following OxyHb treatment (Figure 5D), indicating that OxyHb globally reshapes VGLL3 chromatin occupancy.

To characterize the DNA sequence preferences of VGLL3 binding, we performed motif-enrichment analysis of CUT&Tag peaks. This analysis identified two highly significant motifs (Figure S7): a primary motif with a consensus sequence containing multiple G residues (p = 1e-186) and a secondary motif consisting predominantly of consecutive G bases (p = 1e-179). The enrichment of G-rich motifs is consistent with the GC-rich nature of promoter-proximal regions and supports the preferential binding of VGLL3 to these regulatory elements.

To identify VGLL3-target genes involved in fibrosis, we performed KEGG enrichment analysis of genes adjacent to upregulated peaks, revealing significant enrichment in matrix remodeling-related pathways, including PI3K-Akt signaling, ECM-receptor interaction, focal adhesion, and the TGF-β/Smad pathway (Figure 5E). Genome browser tracks further confirmed VGLL3 occupancy at the *Col1a1* locus (Figure 5F) and other fibrosis-related gene loci (Figures S7C–S7H), including *Tgfb1*, *Col5a1, Plod1*, *Plod2*, *Smad6*, and *Smad7*. ChIP-qPCR validation confirmed VGLL3 occupancy at the promoter regions of these fibrosis-related genes (Figures S7I–S7N).

We next examined the transcriptional consequences of VGLL3 activation by RNA-seq. Volcano plot analysis revealed 202 upregulated and 142 downregulated genes in OxyHb-treated pericytes, among which *Col1a1* exhibited the most pronounced upregulation (Figure 5H). Heatmap analysis further confirmed significant upregulation of fibrosis-associated genes, including *Col1a1*, *Col5a1*, *Col6a1*, *Plod1*, and *Smad6*, following OxyHb stimulation (Figure 5I). RT-qPCR validation confirmed the upregulation of these genes (Figure 5J).

To further investigate the structural basis of VGLL3–DNA recognition, we performed molecular docking and molecular dynamics simulations (Supplemental Figure S8). Consistent with our CUT&Tag and ChIP-qPCR data, the docking models revealed that VGLL3 directly interacts with the *Col1a1* promoter DNA via a conserved binding interface, with key residues, such as Lys-319 and Ser-320, forming stable hydrogen bonds with DNA bases.

Collectively, these findings establish a direct causal link between VGLL3 genomic occupancy and transcriptional activation of fibrogenic genes, particularly *Col1a1,* in pericytes, providing the mechanistic basis for OxyHb-driven pericyte-mediated PFCM formation *in vitro*.

### Pericyte-specific VGLL3 knockout alleviates long-term cerebral autoregulation dysfunction and dilation impairment of the cerebral microcirculation after SAH in mice

Super-resolution ultrasound imaging revealed a progressive reduction in cerebral blood flow velocity and vascular density in *Vgll3^fl/fl^* littermates at 1 and 30 days after SAH. In contrast, these parameters were significantly restored in *Vgll3^ΔPC^* mice (Figures 6A, 6B, and 6D). Photoacoustic imaging further revealed increased cerebrovascular tortuosity in *Vgll3^fl/fl^* littermates at 30 days after SAH, with marked improvement observed in *Vgll3^ΔPC^* mice (Figures 6C and 6E). Consistently, two-photon microscopy revealed preserved vascular architecture in *Vgll3^ΔPC^* mice compared with *Vgll3^fl/fl^* littermates at 30 days after SAH (Figures S9A and S9B).

**Figure 6.**
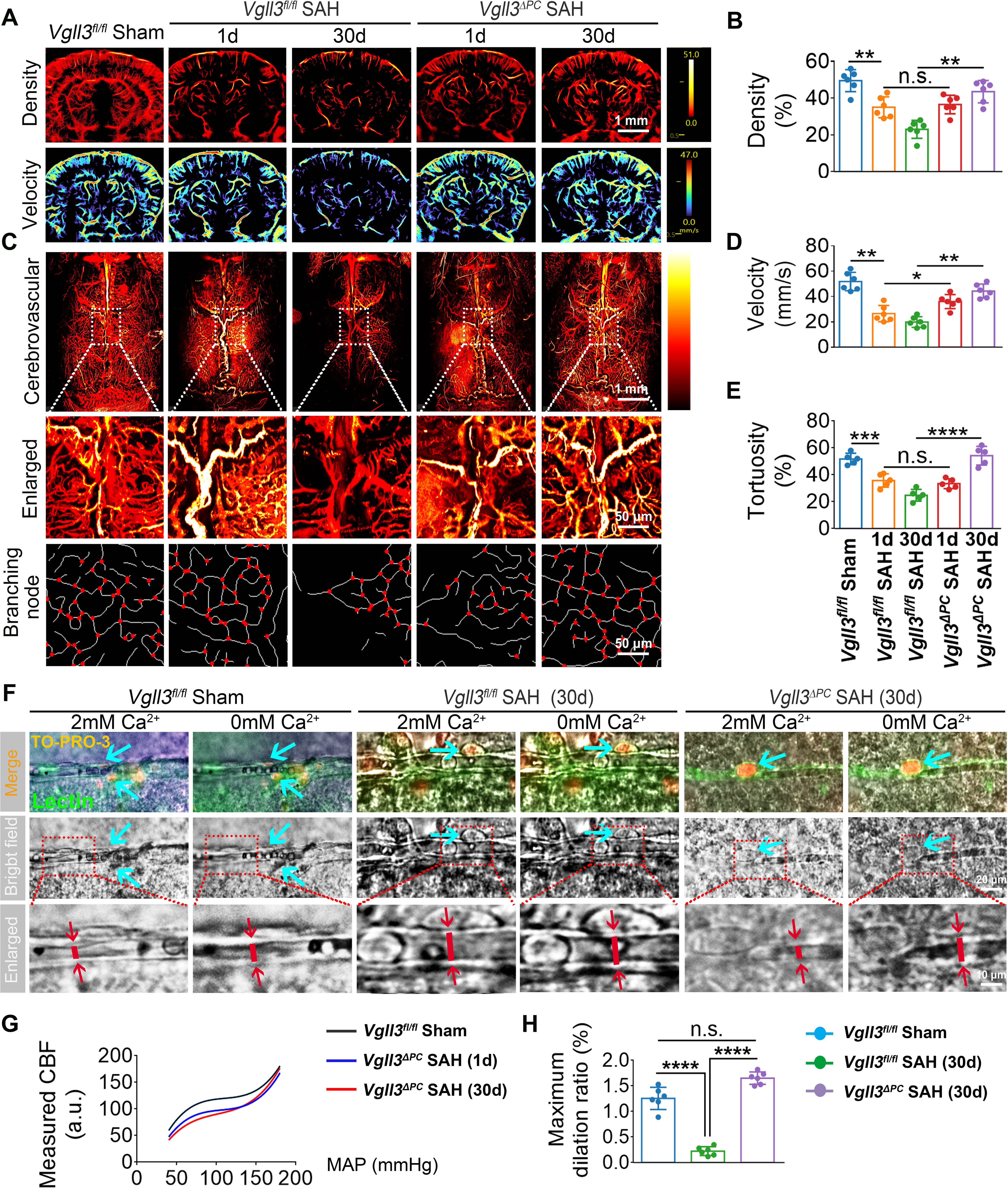
Pericyte-specific VGLL3 deletion improves long-term cerebral autoregulation and microcirculatory dilation after SAH in mice. **A, B**, Cerebral blood flow velocity measured by super-resolution ultrasound imaging in *Vgll3^fl/fl^* littermate and *Vgll3^ΔPC^* mice at 1 and 30 days after SAH. Blood flow velocity progressively declined in *Vgll3^fl/fl^* mice, whereas this reduction was significantly attenuated in *Vgll3^ΔPC^* mice. n = 6 mice per group. Data are presented as mean ± SEM. \*\**p* < 0.01, \*\*\**p* < 0.001 vs. *Vgll3^fl/fl^*; two-way ANOVA followed by Tukey’s post hoc test. **C,** Representative photoacoustic images of cerebrovascular tortuosity in *Vgll3^fl/fl^* and *Vgll3^ΔPC^* littermate mice at 30 days after SAH. Tortuosity was markedly improved in *Vgll3^ΔPC^* mice relative to *Vgll3^fl/fl^* littermates. Scale bar, 50 μm. **D,** Vascular density quantified by super-resolution ultrasound imaging in *Vgll3^fl/fl^* and *Vgll3^ΔPC^* littermate mice at 1 and 30 days after SAH. Vascular density progressively decreased in *Vgll3^fl/fl^* mice, while this decline was significantly attenuated in *Vgll3^ΔPC^* mice. n = 6 mice per group. Data are presented as mean ± SEM. \*\**p* < 0.01, \*\*\**p* < 0.001 vs. *Vgll3^fl/fl^*; two-way ANOVA followed by Tukey’s post hoc test. **E,** Photoacoustic quantification of cerebrovascular tortuosity in *Vgll3^fl/fl^* littermate and *Vgll3^ΔPC^* mice at 30 days after SAH. **F,** Representative images of cerebral microcirculatory dilation during perfusion with Ca²⁺-free ACSF in *Vgll3^fl/fl^* littermate and *Vgll3^ΔPC^* mice at 30 days after SAH. Scale bar, 20 μm. A significant reduction in tortuosity was observed in *Vgll3^ΔPC^* mice compared with *Vgll3^fl/fl^* littermates. **G.** Cerebral autoregulation curves derived from cerebral blood flow velocity and mean arterial pressure (MAP) in in *Vgll3^fl/fl^* sham, *Vgll3^ΔPC^* 1 day, and *Vgll3^ΔPC^* 30 days after SAH. Both *Vgll3^ΔPC^* 1-day and 30-day curves recovered toward the normal range, with only a minimal downward shift compared with *Vgll3^fl/fl^* sham controls. n = 6 mice per group. Data are presented as mean ± SEM. *P < 0.05, \*\**p* < 0.01 vs. *Vgll3^fl/fl^*; unpaired, two-tailed Student’s t-test (tortuosity); two-way ANOVA followed by Tukey’s post hoc test (autoregulation curves). **H,** Maximal vasodilatory response in *Vgll3^fl/fl^* littermate and *Vgll3^ΔPC^* mice at 30 days after SAH. The dilation deficit present in *Vgll3^fl/fl^* mice was markedly reduced in *Vgll3^ΔPC^* mice, especially at pericyte-associated sites. n = 6 mice per group. Data are presented as mean ± SEM. \*\**p* < 0.01 vs. *Vgll3^fl/fl^*; unpaired, two-tailed Student’s t-test. Abbreviations: SAH, subarachnoid hemorrhage; MAP, mean arterial pressure; ACSF, artificial cerebrospinal fluid; SEM, standard error of the mean; n.s., not significant.

We next assessed vasodilatory capacity of the cerebral microcirculation by perfusing with Ca²⁺-free ACSF at 30 days after SAH (Figure 6F). The characteristic dilation impairment observed in *Vgll3^fl/fl^* littermates was markedly alleviated in *Vgll3^ΔPC^* mice, particularly at pericyte-associated sites, with quantitative analysis confirming a significant improvement in maximal vasodilatory response (Figure 6F and 6H). This functional rescue at pericyte locations suggests that pericytic VGLL3 critically regulates dilation capacity of the cerebral microcirculation at the local level.

Cerebral autoregulation curves, generated from cerebral blood flow velocity and mean arterial pressure measurements, revealed that *Vgll3^ΔPC^* mice at both 1 day and 30 days after SAH showed recovery toward the normal range, with only a minimal downward shift compared with *Vgll3^fl/fl^* sham controls (Figure 6G).

Collectively, these findings demonstrate that pericyte-specific VGLL3 knockout not only attenuates PFCM but also rescues long-term cerebral autoregulation dysfunction and cerebral microcirculation dilation impairment after SAH, positioning pericytic VGLL3 as a critical driver of both structural and functional pathology.

## DISCUSSION

The obtained findings determine VGLL3 as the molecular link that couple pericyte hypercontractility to PFCM after SAH, positioning PFCM as the structural basis of long-term cerebral autoregulation dysfunction, especially at lower blood pressure. These findings can extend the mechanistic understanding of DCI beyond the conventional view of self-limiting functional alterations. The heightened susceptibility of SAH patients to cerebral infarction under hypovolemia or hypotension underscores the necessity of rigorous cardiovascular management. Additionally, it offers a rationale for heart-brain integrated therapeutic strategies in SAH survivors.

By using primary cerebral pericytes exposed to OxyHb to model SAH *in vitro*, in combination with *Pdgfrβ^+/-^* models in SAH mice for *in vivo* validation, we established pericytes as the principal cells driving PFCM. Excessive pericyte contraction generates sustained cytoskeletal tension, initiating mechanotransduction cascades^19,21^ that can ultimately drive robust type I collagen deposition in the cerebral microcirculation. Obviously, this pericyte-mediated fibrotic cascade parallels the well-documented perivascular fibrosis in cardiac microcirculation disease,^34^ focusing on a conserved remodeling program across the cardiovascular and cerebrovascular systems. Importantly, unlike reversible acute vasospasm, these collagen-enriched fibrotic lesions are structurally irreversible, offering a mechanistic explanation for why vasospasm-targeted symptomatic treatments fail to resolve persistent neurovascular dysfunction beyond the classic DCI window.

Mechanistically, the present study deals with the core knowledge gap raised in the Introduction: whether VGLL3 transduces pathological pericyte contractile force into profibrotic transcriptional programs eventually compromising long-term cerebral autoregulation after SAH. Considering the lack of publicly available human SAH single-cell transcriptomic data, we initially screened for pericyte-enriched molecules with ICH datasets as a surrogate, considering the shared hemorrhagic pathology and pericyte involvement between the two conditions. Subsequently, the key findings from this screening were validated in our SAH mouse model and *in vitro* OxyHb-treated pericyte system, guaranteeing the translational relevance of our observations. At the molecular level, sustained pericyte contraction is driven by moesin phosphorylation-mediated actin polymerization, promoting the conversion of G-actin to F-actin and generateing cytoskeletal tension. This cytoskeletal tension promotes nuclear translocation of VGLL3, which competes with YAP/TAZ for TEAD binding^35^ to transcriptionally activate collagen-coding genes and drive progressive fibrosis.^24^ Significantly, our data extend this signaling axis to central nervous system pericytes through revealing that VGLL3 directly couples pericyte hypercontractility to PFCM -a novel vasculopathy that underpins long-term cerebral autoregulation dysfunction. Taken together, this VGLL3-driven pathway couples pericyte hypercontractility to irreversible PFCM, causing long-term autoregulatory dysfunction. Moreover, this vasculopathy underpins the progressive neurological deficits consistently found in clinical SAH cohorts,^2,10^ as shown in large-scale outcome studies.

Our findings fundamentally challenge the traditional view, articulated in our Introduction, that SAH-related cerebral ischemia is confined to the acute DCI window. The PFCM found at 30 days after SAH (Figure 1B and 1H) offers a structural foundation for the continued high incidence of cerebral ischemic events documented in long-term cohort studies.^12^ Different from the transient, self-limiting functional alterations of the cerebral microcirculation that have been the traditional focus, PFCM indicates an irreversible structural alteration compromising microcirculation compliance and impairing vasodilatory reserve^36^ — precisely the “novel vasculopathy” that this work defines and that our title emphasizes. This distinction shows deep clinical implications: it indicates that the current clinical focus on acute DCI management neglects a chronic, progressive vasculopathy driving long-term neuropsychiatric deterioration.^37^ Consistent with this notion, the failure of vasospasm-directed interventions to improve DCI outcomes, despite angiographic resolution of vasospasm, points to the existence of a non-vasospastic, structural component of cerebral ischemia, namely the PFCM we have detected.^38,39^ As a result, the obtained findings reframe ischemia after SAH as a biphasic process: an early, reversible functional phase (vasospasm) and a late, irreversible structural phase (PFCM) underlying long-term cerebral autoregulation dysfunction and expanding the boundaries of the conventional DCI window.

Several research biases or confounding factors may have resulted in the ignorance of long-term cerebral ischemia after SAH. First, assessments of cerebral blood flow (CBF) are largely confined to resting-state baseline measurements, neglecting its dynamic characteristics and hemodynamic regulators.^40^ CBF is deeply modulated by blood pressure, arterial blood gases, as well as neuronal signals; therefore, complete hemodynamic profiling requires parallel evaluation of cerebral autoregulation (dependent on blood pressure), cerebrovascular reactivity (dependent on blood gases and vasoactive agents), and neurovascular coupling (dependent on neuronal and sensory activation) .^41^ Besides, neglect of these multidimensional readouts may mask sustained microcirculatory dysfunction persisting long after the acute phase. Secondly, conventional endpoints for evaluating ischemic recovery remain suboptimal. A large number of studies depend on angiographic resolution of large-vessel vasospasm as a surrogate endpoint for ischemic recovery, despite evidence that such angiographic improvement shows no relationship to better functional outcomes.^39^ Nevertheless, this practice is being increasingly challenged. Consistently, clinical evidence has indicated that improvements in angiographic vasospasm do not relate to better functional outcomes.^38,39^ Consistently, emerging clinical data reveals a weak association between angiographic vasospasm and long-term functional outcomes after SAH.^39^ In certain reports, postmortem vasoreactivity assays were carried out on postmortem cerebral vessels; such specimens lose physiological contractile capacity, rendering these measurements inherently unreliable.^42^ Thirdly, iatrogenic cerebral ischemia secondary to aneurysm repair surgery temporally overlaps with the classic DCI window. This overlap can complicate the distinction between perioperative ischemic injury and spontaneous DCI, as a significant proportion of post-SAH infarcts are caused by treatment-related complications rather than vasospasm,^43,44^ potentially overestimating DCI incidence and diverting attention from true long-term ischemic sequelae.^43^

Moreover, our findings can reveal dilation impairment of the cerebral microcirculation within the lower blood pressure range at 30 days after SAH in mice, echoing the finding of progressive long-term neurological deficits in clinical SAH cohorts. Mechanistically, cerebral resistance vessels dilate in response to falling perfusion pressure as a main autoregulatory defense against ischemia. Nevertheless, in hypertension, structural thickening and luminal narrowing of the cerebral microcirculation shift the lower limit of autoregulation to a higher pressure level, rendering the brain vulnerable to hypotension-induced ischemia.^45^ This observation offers a mechanistic rationale for current clinical practices emphasizing avoidance of hypotension in SAH patients,^46^ while extending this rationale through determining PFCM—a structural vasculopathy extending beyond the acute vasospasm phase. Therefore, this study proposes that it is essential for blood pressure management strategies to consider not only acute DCI risk but also long-term structural integrity of the microcirculation. This rigid, collagen-encased microcirculation fails to dilate when perfusion pressure drops, rendering it exquisitely vulnerable to even mild hypotension. Therefore, SAH patients with PFCM may confront elevated risk of cerebral infarction under hypovolemic or hypotensive conditions, highlighting the need for stringent cardiovascular management within a heart-brain integrated therapeutic paradigm.

This study still had the following limitations. At first, our findings are derived exclusively from a mouse SAH model induced by autologous blood injection, which does not completely recapitulate human aneurysmal SAH. Validation in other preclinical models is warranted. Secondly, while we identified VGLL3 as a critical mediator in pericytes, we cannot exclude its potential function in other vascular cell types. Thirdly, the functional consequences of pericyte-specific VGLL3 knockout on normal cerebral microcirculatory homeostasis remain incompletely understood. Finally, even though our data robustly support a causative role for VGLL3 in PFCM, the translational gap is substantial, as small-molecule modulators of VGLL3 are not yet available. Future researches should prioritize (1) inhibiting pericyte hypercontractility to reduce PFCM and assessing blood-brain barrier integrity under long-term ischemic conditions, and (2) developing pharmacologic VGLL3 inhibitors, targeting upstream regulators, as well as establishing non-invasive imaging for early PFCM detection in SAH patients.

To conclude, we have determined VGLL3 as a critical molecular link between pericyte hypercontractility and PFCM, defining a previously unrecognized vasculopathy driving long-term cerebral autoregulation dysfunction after SAH. Through establishing that SAH-induced cerebral ischemia extends beyond the acute DCI window and involves irreversible PFCM, our findings challenge the prevailing paradigm that focuses exclusively on transient vasospasm. Moreover, these insights identify VGLL3 as a promising therapeutic target for preventing long-term ischemic complications. Additionally, our findings also emphasize the clinical reality that the DCI time window coincides with the perioperative period of heightened iatrogenic risk. This confluence has obscured the recognition of chronic microcirculatory pathology. Moreover, through associating pericyte-driven PFCM with a conserved mechanosensitive pathway shared across cardiac and cerebral microcirculatory diseases, this study offers a theoretical foundation for the heart-brain co-therapy paradigm in multi-organ fibrotic microcirculatory disorders.

## Nonstandard Abbreviations and Acronyms

ACSF: artificial cerebrospinal fluid
CBFV: cerebral blood flow velocity
ChIP-qPCR: chromatin immunoprecipitation quantitative polymerase chain reaction
COL-I: type I collagen
DCI: delayed cerebral ischemia
IF: immunofluorescence
MAP: mean arterial pressure
OxyHb: oxygenated hemoglobin
*Pdgfrβ^+/-^*: pericyte-deficient PDGFRβ heterozygous knockout mice
PFCM: perivascular fibrosis of the cerebral microcirculation
SAH: subarachnoid hemorrhage
SEM: standard error of the mean
TEM: transmission electron microscopy
TSS: transcription start site
*Vgll3^fl/fl^*: *_Vgll3_*^flox/flox^
*Vgll3^ΔPC^*: *Vgll3^flox/flox^*; *Pdgfrβ-*CreERT2 (pericyte-specific *Vgll3* conditional knockout mice)
VGLL3: vestigial-like family member 3
WB: Western blotting

## Supplemental Material

Figures S1–S9 Tables S1–S4

## REFERENCES

1. Smith M, Citerio G. What’s new in subarachnoid hemorrhage. Intensive Care Med. 2015;41:123–126. doi: 10.1007/s00134-014-3548-5

2. Aydin S, Peker S. Long-term cognitive decline after subarachnoid hemorrhage: pathophysiology, management, and future directions. Stroke. 2025;56:1106–1111. doi: 10.1161/strokeaha.124.049969

3. Francoeur CL, Mayer SA. Management of delayed cerebral ischemia after subarachnoid hemorrhage. Crit Care. 2016;20:277. doi: 10.1186/s13054-016-1447-6

4. Marques IP, Albuquerque CRC, Souza NVO, Andrade JBC, Silva GS, Kurtz P. Delayed cerebral ischemia after aneurysmal subarachnoid hemorrhage: a narrative review. Arq Neuropsiquiatr. 2025;83:1–14. doi: 10.1055/s-0045-1809885

5. Budohoski KP, Czosnyka M, Smielewski P, Kasprowicz M, Helmy A, Bulters D, Pickard JD, Kirkpatrick PJ. Impairment of cerebral autoregulation predicts delayed cerebral ischemia after subarachnoid hemorrhage: a prospective observational study. Stroke. 2012;43:3230–3237. doi: 10.1161/strokeaha.112.669788

6. Koide M, Ferris HR, Nelson MT, Wellman GC. Impaired cerebral autoregulation after subarachnoid hemorrhage: a quantitative assessment using a mouse model. Front Physiol. 2021;12:688468. doi: 10.3389/fphys.2021.688468

7. Østergaard L, Aamand R, Karabegovic S, Tietze A, Blicher JU, Mikkelsen IK, Iversen NK, Secher N, Engedal TS, Anzabi M, et al. The role of the microcirculation in delayed cerebral ischemia and chronic degenerative changes after subarachnoid hemorrhage. J Cereb Blood Flow Metab. 2013;33:1825–1837. doi: 10.1038/jcbfm.2013.173

8. Tso MK, Macdonald RL. Acute microvascular changes after subarachnoid hemorrhage and transient global cerebral ischemia. Stroke Res Treat. 2013;2013:425281. doi: 10.1155/2013/425281

9. Hoh BL, Ko NU, Amin-Hanjani S, Chou SY, Cruz-Flores S, Dangayach NS, Derdeyn CP, Du R, Hänggi D, Hetts SW, et al. 2023 guideline for the management of patients with aneurysmal subarachnoid hemorrhage: a guideline from the American Heart Association/American Stroke Association. Stroke. 2023;54:e314–e370. doi: 10.1161/str.0000000000000436

10. de Trizio I, Ferrario A, Bögli SY, Casagrande F, Garcia Alzamora M, Sebök M, Bartussek J, Brandi G. Longitudinal trajectories of functional outcome following aneurysmal subarachnoid hemorrhage: a retrospective study. Crit Care. 2026;30:59. doi: 10.1186/s13054-025-05808-7

11. Fernandez-Perez I, Giralt-Steinhauer E, Cuadrado-Godia E, Guimaraens L, Vivas E, Saldaña J, Suárez-Pérez A, Macias-Gomez A, Revert-Barbera A, Estragues-Gazquez I, et al. Long-term vascular events after subarachnoid hemorrhage. J Neurol. 2022;269:6036–6042. doi: 10.1007/s00415-022-11255-z

12. Parasram M, Parikh NS, Merkler AE, Ch’ang JH, Navi BB, Kamel H, Zhang C, Murthy SB. Long-term risk of ischemic stroke among elderly survivors of non-traumatic subarachnoid hemorrhage. Cerebrovasc Dis. 2022;51:14–19. doi: 10.1159/000517416

13. Balbi M, Vega MJ, Lourbopoulos A, Terpolilli NA, Plesnila N. Long-term impairment of neurovascular coupling following experimental subarachnoid hemorrhage. J Cereb Blood Flow Metab. 2020;40:1193–1202. doi: 10.1177/0271678x19863021

14. Han F, Clancy U, Arteaga-Reyes C, Thrippleton MJ, Valdés Hernández MDC, Jaime Garcia D, Stringer MS, Backhouse E, Chappell FM, Cheng Y, et al. Implications of cranial arterial stenosis and dolichoectasia for cerebral small-vessel disease etiopathogenesis: findings from a prospective mild stroke cohort. Circulation. 2026;153:1813–1826. doi: 10.1161/circulationaha.126.079493

15. Jiang Z, Sorrentino G, Simsek S, Roelofs J, Niessen HWM, Krijnen PAJ. Increased perivascular fibrosis and pro-fibrotic cellular transition in intramyocardial blood vessels in myocardial infarction patients. J Mol Cell Cardiol Plus. 2024;10:100275. doi: 10.1016/j.jmccpl.2024.100275

16. Fangma YJ, Zhou ZC, Chen Z, Zheng YR. Translational insights into pericyte-mediated regulation of cerebral blood flow: implications for ischemic stroke. Stroke. 2025;56:e254–e266. doi: 10.1161/strokeaha.125.052018

17. Li Y, Zhou L, Deng H, Zhang Y, Li G, Yu H, Wu K, Wang F. A switch in the pathway of TRPC3-mediated calcium influx into brain pericytes contributes to capillary spasms after subarachnoid hemorrhage. Neurotherapeutics. 2024;21:e00380. doi: 10.1016/j.neurot.2024.e00380

18. Dalkara T, Østergaard L, Heusch G, Attwell D. Pericytes in the brain and heart: functional roles and response to ischaemia and reperfusion. Cardiovasc Res. 2025;120:2336–2348. doi: 10.1093/cvr/cvae147

19. Dessalles CA, Babataheri A, Barakat AI. Pericyte mechanics and mechanobiology. J Cell Sci. 2021;134. doi: 10.1242/jcs.240226

20. Wilhelm I, Győri F, Dudás T, Nagy V, Shreeya T, Krecsmarik M, Farkas AE, Fazakas C, Krizbai IA. Eppur si muove: the dynamic brain pericyte. Fluids Barriers CNS. 2025;22:95. doi: 10.1186/s12987-025-00706-0

21. Kotecki M, Zeiger AS, Van Vliet KJ, Herman IM. Calpain-and talin-dependent control of microvascular pericyte contractility and cellular stiffness. Microvasc Res. 2010;80:339–348. doi: 10.1016/j.mvr.2010.07.012

22. Kureli G, Yilmaz-Ozcan S, Erdener SE, Donmez-Demir B, Yemisci M, Karatas H, Dalkara T. F-actin polymerization contributes to pericyte contractility in retinal capillaries. Exp Neurol. 2020;332:113392. doi: 10.1016/j.expneurol.2020.113392

23. Rolle IG, Crivellari I, Zanello A, Mazzega E, Dalla E, Bulfoni M, Avolio E, Battistella A, Lazzarino M, Cellot A, et al. Heart failure impairs the mechanotransduction properties of human cardiac pericytes. J Mol Cell Cardiol. 2021;151:15–30. doi: 10.1016/j.yjmcc.2020.10.016

24. Horii Y, Matsuda S, Toyota C, Morinaga T, Nakaya T, Tsuchiya S, Ohmuraya M, Hironaka T, Yoshiki R, Kasai K, et al. VGLL3 is a mechanosensitive protein that promotes cardiac fibrosis through liquid-liquid phase separation. Nat Commun. 2023;14:550. doi: 10.1038/s41467-023-36189-6

25. Liu Y, Yang Z, Lin N, Liu Y, Chen H. Highly expressed VGLL3 in keloid fibroblasts promotes glycolysis and collagen production via the activation of Wnt/β-catenin signaling. Cell Signal. 2025;127:111604. doi: 10.1016/j.cellsig.2025.111604

26. Kaminski WE, Lindahl P, Lin NL, Broudy VC, Crosby JR, Hellström M, Swolin B, Bowen-Pope DF, Martin PJ, Ross R, et al. Basis of hematopoietic defects in platelet-derived growth factor (PDGF)-B and PDGF beta-receptor null mice. Blood. 2001;97:1990–1998. doi: 10.1182/blood.v97.7.1990

27. Deng HJ, Deji Q, Zhaba W, Liu JQ, Gao SQ, Han YL, Zhou ML, Wang CX. A20 establishes negative feedback with TRAF6/NF-κB and attenuates early brain injury after experimental subarachnoid hemorrhage. Front Immunol. 2021;12:623256. doi: 10.3389/fimmu.2021.623256

28. Zhang YJ, Li YC, Yu HF, Li C, Deng HJ, Dong YH, Li GB, Wang F. Imaging vital and non-vital brain pericytes in brain slices following subarachnoid hemorrhage. J Vis Exp. 2023. doi: 10.3791/65873

29. Balbi M, Ghosh M, Longden TA, Jativa Vega M, Gesierich B, Hellal F, Lourbopoulos A, Nelson MT, Plesnila N. Dysfunction of mouse cerebral arteries during early aging. J Cereb Blood Flow Metab. 2015;35:1445–1453. doi: 10.1038/jcbfm.2015.107

30. Lunde IG, Rypdal KB, Van Linthout S, Diez J, González A. Myocardial fibrosis from the perspective of the extracellular matrix: Mechanisms to clinical impact. Matrix Biol. 2024;134:1–22. doi: 10.1016/j.matbio.2024.08.008

31. Pennings FA, Albrecht KW, Muizelaar JP, Schuurman PR, Bouma GJ. Abnormal responses of the human cerebral microcirculation to papaverin during aneurysm surgery. Stroke. 2009;40:317–320. doi: 10.1161/strokeaha.108.522375

32. Karvar S, Ansa-Addo EA, Suda J, Singh S, Zhu L, Li Z, Rockey DC. Moesin, an ezrin/radixin/moesin family member, regulates hepatic fibrosis. Hepatology. 2020;72:1073–1084. doi: 10.1002/hep.31078

33. Cuervo H, Pereira B, Nadeem T, Lin M, Lee F, Kitajewski J, Lin CS. PDGFRβ-P2A-CreER(T2) mice: a genetic tool to target pericytes in angiogenesis. Angiogenesis. 2017;20:655–662. doi: 10.1007/s10456-017-9570-9

34. Tamiato A, Tombor LS, Fischer A, Muhly-Reinholz M, Vanicek LR, Toğru BN, Neitz J, Glaser SF, Merten M, Rodriguez Morales D, et al. Age-dependent RGS5 loss in pericytes induces cardiac dysfunction and fibrosis. Circ Res. 2024;134:1240–1255. doi: 10.1161/circresaha.123.324183

35. Ma S, Tang T, Probst G, Konradi A, Jin C, Li F, Gutkind JS, Fu XD, Guan KL. Transcriptional repression of estrogen receptor alpha by YAP reveals the Hippo pathway as therapeutic target for ER(+) breast cancer. Nat Commun. 2022;13:1061. doi: 10.1038/s41467-022-28691-0

36. Lu S, Strand KA, Mutryn MF, Tucker RM, Jolly AJ, Furgeson SB, Moulton KS, Nemenoff RA, Weiser-Evans MCM. PTEN (phosphatase and tensin homolog) protects against Ang II (angiotensin II)-induced pathological vascular fibrosis and remodeling-brief report. Arterioscler Thromb Vasc Biol. 2020;40:394–403. doi: 10.1161/atvbaha.119.313757

37. Diestro JDB, Vyas M, Jung Y, Kishibe T, Leochico C, Espiritu A, Dorotan MK, Dimal N, Omar AT, Sienes A, et al. Long-term neuropsychiatric complications of aneurysmal subarachnoid hemorrhage: a narrative review. J Neurointerv Surg. 2025;17:167–173. doi: 10.1136/jnis-2023-020979

38. Etminan N, Vergouwen MD, Macdonald RL. Angiographic vasospasm versus cerebral infarction as outcome measures after aneurysmal subarachnoid hemorrhage. Acta Neurochir Suppl. 2013;115:33–40. doi: 10.1007/978-3-7091-1192-5_8

39. Behzadi F, Tsiang JT, Jani RH, Payman AA, Bond BJ, Kam AW, Pasquale DD, Serrone JC. Angiographic response to endovascular treatment of post-hemorrhage cerebral vasospasm is not associated with clinical outcome. Clin Neurol Neurosurg. 2025;254:108927. doi: 10.1016/j.clineuro.2025.108927

40. Claassen JA. Shifting concepts of autoregulation: commentary to ‘Static autoregulation in humans’. J Cereb Blood Flow Metab. 2024;44:1397–1399. doi: 10.1177/0271678x241254676

41. Claassen J, Thijssen DHJ, Panerai RB, Faraci FM. Regulation of cerebral blood flow in humans: physiology and clinical implications of autoregulation. Physiol Rev. 2021;101:1487–1559. doi: 10.1152/physrev.00022.2020

42. Onoue H, Kaito N, Akiyama M, Tomii M, Tokudome S, Abe T. Altered reactivity of human cerebral arteries after subarachnoid hemorrhage. J Neurosurg. 1995;83:510–515. doi: 10.3171/jns.1995.83.3.0510

43. Wagner M, Steinbeis P, Güresir E, Hattingen E, du Mesnil de Rochemont R, Weidauer S, Berkefeld J. Beyond delayed cerebral vasospasm: infarct patterns in patients with subarachnoid hemorrhage. Clin Neuroradiol. 2013;23:87–95. doi: 10.1007/s00062-012-0166-x

44. Hu P, Yan T, Li Y, Guo G, Gao X, Su Z, Du S, Jin R, Tao J, Yuan Y, et al. Effect of surgical clipping versus endovascular coiling on the incidence of delayed cerebral ischemia in patients with aneurysmal subarachnoid hemorrhage: a multicenter observational cohort study with propensity score matching. World Neurosurg. 2023;172:e378–e388. doi: 10.1016/j.wneu.2023.01.032

45. Barry DI. Cerebral blood flow in hypertension. J Cardiovasc Pharmacol. 1985;7 Suppl 2:S94–98. doi: 10.1097/00005344-198507002-00018

46. Diringer MN, Bleck TP, Claude Hemphill J, 3rd, Menon D, Shutter L, Vespa P, Bruder N, Connolly ES, Jr., Citerio G, Gress D, et al. Critical care management of patients following aneurysmal subarachnoid hemorrhage: recommendations from the Neurocritical Care Society’s Multidisciplinary Consensus Conference. Neurocrit Care. 2011;15:211–240. doi: 10.1007/s12028-011-9605-9

